# Community stability increases the predictability of microeukaryote community coalescence outcomes

**DOI:** 10.1101/2024.02.14.580275

**Authors:** Máté Vass, Anna Székely, Ulla Carlsson-Graner, Johan Wikner, Agneta Andersson

## Abstract

Mixing of entire microbial communities represents a frequent, yet understudied phenomenon. Here, we mimicked estuarine condition in a microcosm experiment by mixing a freshwater river community with a brackish sea community and assessed the effects of both environmental and community coalescences induced by varying mixing processes on microeukaryotic communities. Signs of shifted community composition of coalesced communities towards the sea parent community suggest asymmetrical community coalescence outcome, which, in addition, was generally less impacted by environmental coalescence. Diatoms were negatively impacted by coalescence, while fungi, ciliates, and cercozoans were promoted to varying extents, depending on the mixing ratios of the source (i.e., river or sea) communities. Community stability, inferred from community cohesion, suggests that the more stable parent community (i.e., community with greater negative cohesion attributed to competitive interactions) dominates the final, coalesced community, but the fate of its community members is influenced by mixing ratios and frequencies (i.e., one-time versus repeated coalescence). Generally, community coalescence increased alpha diversity and promoted competition from the introduction (or emergence) of additional (or rare) species. These competitive interactions in turn had community stabilizing effect as evidenced by the increased proportion of negative cohesion. Our study suggests that the predictability of coalescence outcomes was greater when the more stable parent community (i.e., sea microbes) dominated the final community and this predictability was further enhanced when communities collided repeatedly.

**Open research statement:** Sequencing data is deposited to NCBI SRA database under the accession number PRJNA922225. Data (OTU table, consensus taxonomy and metadata) are available in Open Science Framework (OSF) (http://osf.io/sme36).

## Introduction

Community coalescence is a complex phenomenon that involves the mixing of microbial communities from previously isolated environments (Rillig et al. 2015). Such mixing events comprise the movement and potential mixing of environments resulting in environmental coalescence, as well as the dispersal of microbial communities termed as biotic coalescence. Coalescence events are expected to impose dual impacts on communities in the form of *environmental filtering* due to the altered environment following mixing, and *community reorganization* by the dynamic rearrangement of ecological interactions covering competitive/facilitative interactions, species co-recruitment and trophic interactions (Rillig et al. 2015, Custer et al. 2023). Interestingly, colliding communities maintain some of these biotic interactions as intact, thus, stabilising the coalesced community (also known as network coherence) (Rillig et al. 2015). Hence, the likelihood of community establishment following coalescence can depend on biotic interaction types and the cohesiveness of taxa within the parent communities. The recently developed ‘cohesion’ metric (Herren and McMahon 2017) quantifies the degree to which members of a community are connected and can be utilized to evaluate its impact on community stability (see e.g., Hernandez et al. 2021). Simply, because communities with greater fraction of negative cohesion, attributed to competition and/or environmental filtering, tend to be more stable (e.g., have low species turnover over time) (Herren and McMahon 2017, Bier et al. 2022). In contrast, higher fraction of positive cohesion indicates greater environmental synchrony and/or facilitative interactions between taxa, which in turn results in a community with the potential for mutual downfall (Coyte et al. 2015). The latter happens, for example, when the decreasing abundance of one species pulls others down and by doing so, destabilise the community via positive-feedback loops. Inferring community stability from community coherence, thus, represents a promising avenue for understanding and predicting the outcome of community coalescence. In this case, members of more stable parent community (community with greater negative cohesion, thus, being competition-driven) are expected to dominate the final mixed community (Coyte et al. 2015, Herren and McMahon 2017, Castledine et al. 2020). However, a previous work suggests that parent communities with more facilitative interactions contribute to a greater proportion of species in the final coalesced community due to their superior ability to deplete resources and resist invasions (Chang et al. 2021, Lechón-Alonso et al. 2021), especially when coalescence happens repeatedly (Lechón-Alonso et al. 2021, Song et al. 2021). Thus, the temporal scale in which coalescence events occur, for example, the frequency of invasion events (i.e., one-time versus repeated coalescence) have most likely a key role in determining the dominance of interaction type of a community network, and consequently, coalescence outcomes.

Coalescence events are particularly common in aquatic ecosystems as water bodies of different origin often mix at interfaces like river–sea junctions (Rillig and Mansour 2017). Such estuary habitats present challenging environments for microbial communities in respect of salinity, oxygen levels, and nutrient concentrations, which vary greatly not only spatially along the mixing zones, but also temporarily as a result of the fluctuations that emerge due to hydrological features of river inflows and tidal intrusions (Wolanski et al. 2012, Lee et al. 2017, Mansour et al. 2018). Such spatio-temporal environmental variability drives the development of diverse microbial communities, often characterised by protistan species maxima (Telesh et al. 2011). In estuaries, communities continuously coalesce and form a new set of populations with different community structure and stability. Only microbes that are able to adjust their osmoregulation and metabolic profiles (i.e., nutrient acquisition) and/or elevate their growth rates can survive these rapidly changing conditions (Bouvier and del Giorgio 2002, Balzano et al. 2015, Tee et al. 2021). Such river-sea mixing events are particularly common in the Baltic Sea which is a shallow brackish sea characterized by large river influence (Raudsepp et al. 2023) making it a perfect environment to study the process of whole-community mixing. Past works suggest that even the low salinity of the Baltic Sea impacts river-transported microbes lacking adaptability to saline conditions (Langenheder et al. 2003, Shen et al. 2018), shifting the final, mixed community towards that of the sea (Székely et al. 2013, Rocca et al. 2020, Song et al. 2022). However, evidence for riverine prokaryotic species (*Flavobacteria* spp. and *Marinomonas* sp.) immigrating and growing in the brackish water environment has also been reported from the northern Baltic Sea (Kisand et al. 2005).

Although microeukaryotes play crucial roles in aquatic primary production and nutrient cycling via their roles in the microbial loop, only a few studies aimed to investigate the eukaryotic fraction of the microbial consortia along river-to-sea transects (Yang et al. 2021, Tee et al. 2021, Vass et al. 2022). Community coalescence and the mechanisms underlying its outcomes are nevertheless scarcely studied along river to sea transitions (Mansour et al. 2018). Therefore, we aimed to mimic estuarine condition in a microcosm experiment by mixing freshwater river community with brackish sea community and specifically investigate whether the fate of microeukaryotes during community coalescence differ in response to varying mixing frequencies and ratios.

Here we address two fundamental questions about community coalescence: (i) Does mixing ratio define the community coalescence outcomes? (ii) What effect does mixing frequency (one-time versus repeated coalescence) have on the final community composition? (iii) What is the role of community stability during community coalescence? Beginning with two microbial communities originating from a river and an offshore site in the Gulf of Bothnia, we inoculated each of them separately in their mixed environment to assess environmental coalescence. Community coalescence outcomes were also assessed after mixing them one-time or repeatedly at three different mixing ratios, varying the initial ratio of the parent communities. We hypothesized that (a) it is possible to predict the outcome of community coalescence based on the applied mixing ratios and the individual environmental adaptive capabilities of the parent communities, and further, (b) that one-time coalescence is advantageous for competition-driven (stable) communities, while repeated mixing of communities eventually results in facilitation-dominated communities, as suggested by Lechón-Alonso et al. (2021).

## Material and methods

### Sampling

The two microbiomes for our microcosm experiment were collected during the diatom spring bloom on April 25, 2022, from a coastal area of the Gulf of Bothnia, Sweden. The freshwater river sample originated from a coastal river, Ängerån (63°34’51.8“N; 19°50’07.0”E). The anthropogenic effect (i.e., pollution) on this river is negligible since it mostly runs through coniferous forests and over relatively young marine sediment (Müller 1982). The brackish sea water was collected from an offshore site (63°28’30.20”N; 19°50’5.85”E), approximately 12 km from the mouth of the river Ängerån. River and sea samples were taken from the euphotic zone (integrated sample from 0-10 m depth, in the case of sea water) and transported to the laboratory in sterile containers. The samples were pre-filtered through a 200 μm mesh to remove macroorganisms (i.e., mesozooplankton) and debris, and used immediately to set up the experiment. Total dissolved nitrogen (TDN) and phosphorus (TDP) were also measured, following standard analytical methods described in (Hansen and Koroleff 1999).

### Coalescence experiment

A 16-day long experiment was conducted, using parent communities prepared from the river (R) and sea (S) samples (Fig. 1). These parent communities (R and S) were then exposed to one-time (OC) or repeated (RC) coalescences in three river:sea mixing ratios (1:1, 1:2 and 2:1).

**Figure 1.**
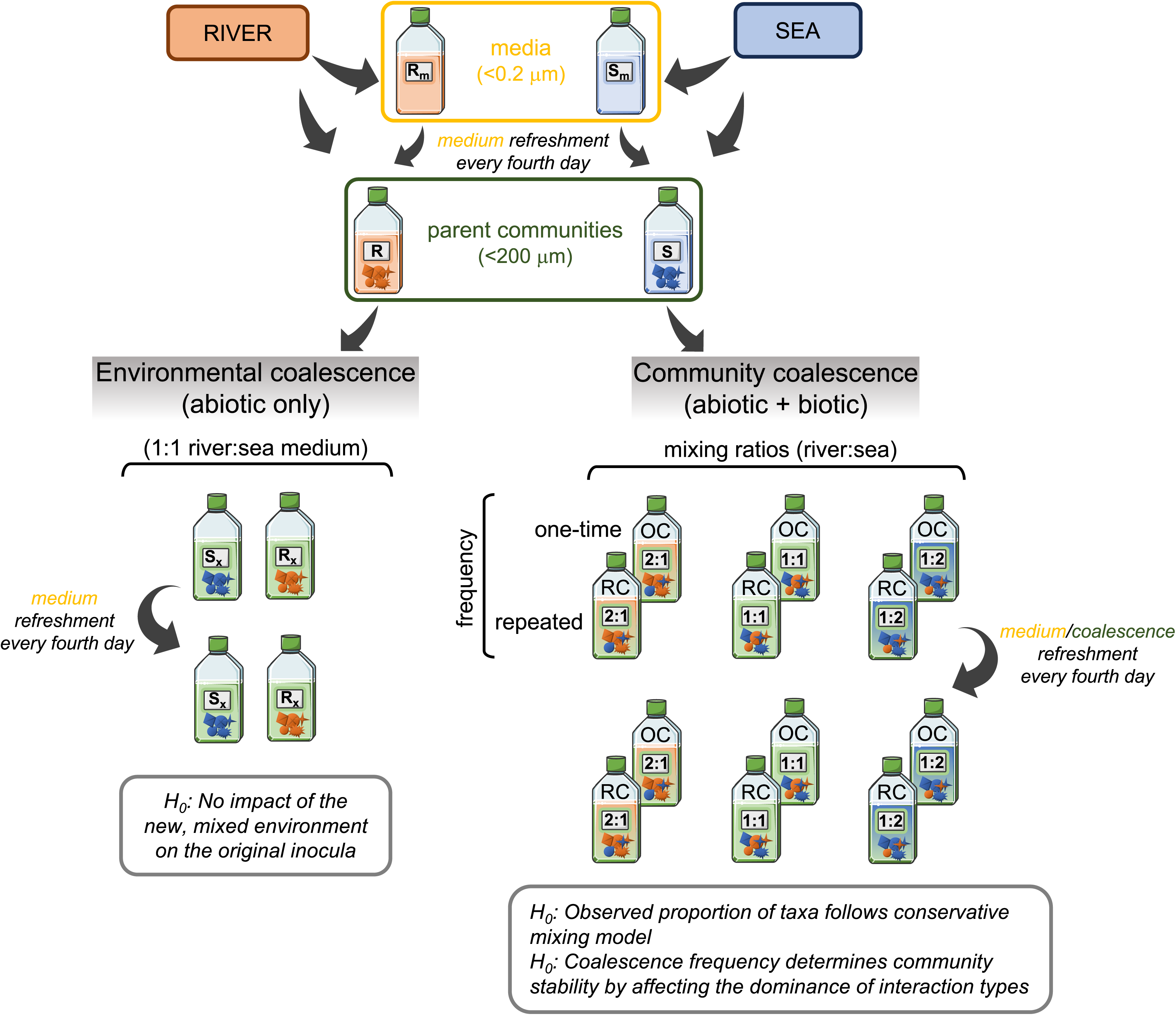
Overview of the experimental design. Pre-filtered (<200 μm) water samples from two sites (river and sea) were collected to inoculate our microcosms. *Environmental coalescence*: each parent community (R, S) was used to assess the potential environmental filtering effect of the mixed medium (river:sea = 1:1) on the unmixed parent communities (R_x_and S_x_). *Community coalescence*: two different coalescence treatments (one-time – OC, and repeated – RC) were set up as follows. River and sea parent communities were mixed at three mixing ratios (1:1, 1:2 and 2:1, resp.) to create coalesced cultures. For the cultures exposed to environmental coalescence and the ones undergoing OC treatment, sterile-filtered (<0.2 μm) media (R_m_ and S_m_) were used during the course of the experiment (i.e., every fourth day) to refresh the parent communities. Cultures exposed to repeated coalescence (RC) received both parent communities mixed according to the corresponding coalescence treatment instead of the filtered media. All experimental treatments (each of 60 mL) consisted of five replicates.

To avoid substantial chemical changes due to autoclaving, the cell-free river (R_m_) and sea (S_m_) media was prepared by sequentially filtration through GF/F filters (0.7 μm, Whatman) and then through sterile 0.2 μm 47 mm membrane filters (Pall Supor) in a laminar flow hood.

To assess the individual environmental adaptive capabilities of the parent communities and estimate the effect of coalescence imposed solely by the abiotic environmental changes of the mixed media (i.e., environmental coalescence), river and sea inoculum (80%; v/v) were incubated separately in blended native media (1:1 mixture of R_m_ + S_m_), hereafter R_x_ and S_x_ (i.e., environmental coalescence), respectively. For the community coalescence treatments, the mixture of R_m_ and S_m_ were inoculated with river (R) and sea (S) parent communities (80%; v/v), according to the applied mixing ratios (e.g., OC_1:2_/RC_1:2_ treatment consisted of 4 mL R_m_ + 8 mL S_m_ and 16 mL R_i_ + 32 mL S_i_) in order to achieve equally-dominated, sea-dominated or river-dominated conditions.

Every four days 20 % sample volume of each OC microcosm was exchanged with the respective medium, following the initial mixing ratios. For this, each replicate ‘A’ of the communities received medium from replicate ‘A’. Likewise, each replicate ‘B’ received medium from replicate ‘B’, etc. Microcosms of RC treatment was exchanged with samples from the respective inoculum communities, instead of the filtered media, applying 20% (v/v) exchange. Both the community coalescence and the medium replacement were carried out in a laminar flow hood, using sterile disposable pipettes.

All treatments (60 mL each) with five replicates were maintained in sterile culture flasks with filter caps (Sarstedt, Nümbrecht, Germany), resulting in 60 cultures in total. The incubation was carried out at 10 °C with a photoperiod set to 17:7 h light:dark cycle to mimic ambient conditions (Andersson et al. 1994). Twice a day, the microcosms were mixed by gently shaking and randomly placed to minimize the potential biases of the differential light in the experiment room.

### Monitoring microcosms during the experiment

Every four days subsamples (12 mL) from each culture flask were pipetted into sterile 15 mL tubes before the exchange of medium and/or parent community and processed as follows. To assess and compare the growth of algae across microcosms, the subsamples were dark-adapted for at least 20 minutes and chlorophyll fluorescence-induced dynamic curve was measured using AquaPen-C device (Photon Systems Instruments, Brno, Czechia). The strong correlation between the integral area of OJIP curve and chlorophyll-a content allows us to estimate chlorophyll content of samples in a quick, non-invasive way (Chen et al. 2021). The validity of this method was checked by measuring the ethanol-extracted (95%) chlorophyll-a concentration of the initial water samples using spectrofluorometer and correlated it with the integral area of the measured OJIP curves (R^2^ = 0.91). We acknowledge that this method has limitations, nevertheless, can be used to monitor our microcosms and to compare them among treatments. After the measurement with AquaPen-C device, subsamples were filtered through 0.2 μm syringe filters and kept frozen until the measurement of chemical properties (e.g., total dissolved nutrients). Total dissolved nitrogen (TDN) and phosphorus (TDP), were measured, following standard analytical methods described in Hansen and Koroleff (1999). At the end of the experiment (day 16) the cultures were filtered by vacuum filtration onto 0.2 μm 47 mm membrane filters (Pall Supor) and the filters were stored at –80 °C.

### Community analysis by long-read amplicon sequencing

The DNA was extracted from the filters using the ZymoBIOMICS DNA Miniprep Kit (Zymo Research Corp, CA, USA) following manufacturer’s protocol. DNA extracts were quantified with NanoDrop (ND-1000 Spectrophotometer).

Amplification was done using the V4_Balzano_F/D11_3143R primer pair (see, Appendix S1: Table S1) in order to amplify almost the whole (∼4.5 kbp) eukaryotic rRNA operon (Latz et al. 2022), which allows better taxonomic classification. The PCR was performed according to (Latz et al. 2022), using 20 ng template DNA. The barcoded PCR products were purified with 0.8× of AMPure magnetic beads (Beckmann) following the manufacturer’s protocol. Thereafter, the purified PCR products were quantified using the Qubit 1× HS Assay Kit (ThermoFisher Scientific) and pooled in equimolar amounts.

One µg of library was used for the ONT library preparation using the 1D sequencing (SQK-LSK109; Oxford Nanopore Technologies), following some modifications described in Vass et al. (2022). Sequencing was performed using a MinION Mk1C instrument (ONT) operated with a Spot-ON Flow Cell (R9.4.1 chemistry). Real-time high-accuracy basecalling (HAC) was executed using the MinKNOW software (v22.05.6), resulting in 2.43 M reads Q > 9.

Quality reads were demultiplexed and barcoded primers were trimmed with MiniBar (Krehenwinkel et al. 2019), filtered by length (2–6 kb) with NanoFilt (v2.8.0) (De Coster et al. 2018), and processed using the NGSpeciesID pipeline (v0.1.2.2) (Sahlin et al. 2021, Pomerantz et al. 2022) with –mapped_threshold 0.8 –aligned_threshold 0.6 parameters during read clustering by isONclust (Sahlin and Medvedev 2020). We obtained on average 29,078 reads per sample with 62.1% mapping rate (i.e. percent of high-quality paired reads for generation of the consensus sequences from total reads) (Appendix S1: Table S2). Quality-filtered (Q > 9 and 2–6 kb) reads were deposited to NCBI SRA database under the accession number PRJNA922225.

18S, 28S rRNA genes (SSU and LSU) and the full length internal transcribed spacer (ITS) were extracted using ITSx (Bengtsson-Palme et al. 2013) and used in BLASTn search to assign taxonomy against the PR2 v4.14 database (Guillou et al. 2013), SILVA LSU v138.1 reference database (Quast et al. 2012) and the UNITE+INSD v9.0 database (Abarenkov et al. 2022), respectively, using BLAST+ (v2.11.0+) and keeping hits with at least 80 % identity. Results of the BLASTn search were processed with phyloR (https://github.com/cparsania/phyloR) to keep top hits and to assign taxonomy levels. Each OTU was manually inspected, and consensus classification was determined down only to the level that could be robustly supported by at least two of the three reference databases, using the 2 out of 3 rule (e.g., if an OTU classified as taxon A by two reference databases but as taxon B by the third one, then taxon A is selected). Unidentified (non-eukaryotic) consensus sequences and their corresponding OTUs from the OTU table were discarded (n = 130). The taxonomic distribution of reads was visualized with Krona (http://sourceforge.net/projects/krona).

### Data analysis

All statistical analyses and visualizations were conducted in R version 4.0.4 (R Core Team 2021). Rarefaction was done using 4,752 reads per sample, resulting in 461 OTUs. The final OTU table and the corresponding taxonomy are available in Open Science Framework (OSF) (https://osf.io/sme36). Sample coverage was assessed with the ‘iNext’ R package (Hsieh et al. 2016), and found that community composition was sufficiently covered (Appendix S2: Figure S1). Differences in total dissolved nutrients (i.e., TDN and TDP) across inoculum sources and treatments were assessed by pairwise Wilcoxon rank-sum test with a Benjamini-Hochberg (BH) corrected significance cutoff of 0.05. Diversity analyses (alpha-diversity and beta-diversity based on Bray–Curtis distance) were performed using the ‘microeco’ R package (v.0.6.5) (Liu et al. 2021) and the results (i.e. nonmetric multidimensional scaling – NMDS) were plotted using ‘ggplot’ package (Wickham 2009). Difference in alpha diversity across inoculum sources and treatments were tested with ANOVA followed by Duncan’s test (*p* < 0.05) as a post-hoc test. To test compositional differences between samples, pairwise permutational multivariate analysis of variance (PERMANOVA, permutations: 999) was performed using the function pairwise.adonis in ‘pairwiseAdonis’ R package (Arbizu 2019).

### Community stability inferred from community cohesion

To estimate network coherence that impacts community stability, we first quantified the abundance-weighted pairwise correlations of every OTU and used the resulting positive and negative co-occurrences separately to calculate negative and positive community cohesion, respectively, as proposed by Herren and McMahon (2017). Cohesion is a metric that measures the degree of connectivity of each observed microbial community. Throughout this paper, we infer community stability from the absolute value of the ratio of negative and positive cohesion (negative:positive), as in Hernandez et al. (2021). This community stability metric takes < |1| values when communities have higher proportions of facilitation than competition, while values > |1| suggest competition-dominated communities, and thus, a community with more negative-feedback loops. Through such negative-feedback loops propagation of perturbations to the rest of the community is dampened, leading to greater overall community stability (Fontaine et al. 2011, Coyte et al. 2015).

Note that for the estimation of community stability, we used non-rarefied dataset as suggested by Herren and McMahon (2017). Differences in community stability across inoculum sources and treatments were assessed by pairwise Wilcoxon rank-sum test with a Benjamini-Hochberg (BH) corrected significance cutoff of 0.05.

### Evaluation of community coalescence

Changes in the relative abundance of each OTU in the parent inoculum communities (i.e., R or S) compared to their abundance in mixed media of the environmental coalescence treatment (R_x_ or S_x_) were assessed by differential abundance analysis using ZicoSeq (permutation: 999) (Yang and Chen 2022). Only taxa that were present in both the parent community (i.e, R or S) and the corresponding environmental coalescence treatments (i.e., R_x_ or S_x_), and were not affected by the effect of environmental coalescence (i.e., showed no significant (*p*_FDR.adj_ < 0.05) decrease in abundance in R_x_/S_x_ compared to R/S) were selected for further analyses. This ensured to filter out taxa affected by environmental (abiotic) coalescence (i.e., due to environmental filtering) and allowed us to assess the biotic component of community coalescence.

To evaluate the outcome of such community coalescence (i.e. biotic component of community coalescence), we quantified the extent of deviation between the observed coalesced communities and those expected according to a conservative mixing model as in Székely & Langenheder (2017) and Vass et al. (2021) (but see detailed calculation in Appendix S2: Equation S1).

Thereafter, we calculated the Bray-Curtis similarities of the observed and expected coalesced communities. The comparison of the Bray-Curtis similarities was used to indicate the predictability of coalescence outcomes and tested using t-tests. Specifically, no significant deviation (*p* < 0.05) indicates that the observed community does not differ significantly from the expected one, suggesting predictable community coalescence. Differences between community coalescence treatments were assessed using a one-way ANOVA and a subsequent Tukey’s HSD test. Additionally, Bray-Curtis similarities between coalesced communities (i.e., OC and RC) and the corresponding parent communities (i.e., R and S) for the observed and expected data matrices were also calculated, separately. Here, a significantly greater deviation between observed versus expected similarity (*p* < 0.05) indicates that coalescence resulted in greater community divergence from the parent communities than expected. On the other hand, a significantly lower deviation (*p* < 0.05) indicates a higher convergence towards the parent communities than expected, which could be a consequence of asymmetric coalescence outcome (i.e., the dominance of one parent community in the final, coalesced community).

To assess population dynamics in response to community coalescence, we used further differential abundance analyses (ZicoSeq; permutation: 999) to detect OTUs with significant (*p*_FDR.adj_ < 0.05) increase or decrease in taxa abundances in the coalesced communities compared to their abundances in the parent communities (R, S). Finally, Kruskal-Wallis test (since the assumptions of two-way ANOVA were not met) were applied to reveal whether the different coalescence treatments, or mixing ratios, resulted in different total relative abundance of OTUs that increased or decreased after community coalescence.

## Results

### Environmental condition of microcosms

Our initial parent (R, S) and coalesced communities (OC_1:1/1:2/2:1_ and RC_1:1/1:2/2:1_) showed distinct chemical and compositional properties. Total dissolved nutrients (i.e., TDN and TDP), as well as chlorophyll-a concentration – as a proxy of the biomass of primary producers – showed variation across microcosms (Appendix S2: Figure S2, S3). In general, sea medium (S_m_) was nitrogen-poor compared to the river medium (R_m_) (*p* < 0.05), and both had low levels of dissolved phosphorus (TDP). Microcosms exposed to abiotic coalescence (i.e., R_x_, S_x_) did not suggest nutrient-poor conditions by the end of the experiment, as they showed higher or similar TDP and TDN values than those observed in filtered media (S_m_, R_m_) (Appendix S2: Figure S2). Cultures of community coalescence treatments had even greater (Kruskal-Wallis: *p_adj_* < 0.05) availability of TDP than all the other microcosms (except sea inoculum), and their TDN levels were intermediate between the levels of R_m_ and S_m_, showing significant differences (Kruskal-Wallis: *p_adj_* < 0.05) in relation to the applied mixing ratios. The total dissolved nutrients, however, did not differ between one-time and repeated coalescence treatments.

Our inocula originated from oligotrophic ecosystems, hence, the overall observed low values of chlorophyll-a are not peculiar. Estimated biomass of primary producers was significantly higher (Tukey’s HSD: *p* < 0.001) in sea (S) than in river (R) parent communities (Appendix S2: Figure S3). By the end of the experiment (i.e., day 16), these parent communities also significantly differed (Tukey’s HSD: *p* < 0.001) from their respective communities that have been exposed to mixed media (S_i_ vs S_x_ and R_i_ vs R_x_). Interestingly, microcosms with sea inoculum reached higher chlorophyll-a levels when incubated in the mixed (S_x_) than in their original medium (S). In contrast, river community grew better in their original medium (R) compared to the mixed environment (R_x_). Biomass values in both coalescence frequency treatments converged by day 16 (Tukey’s HSD: *p* > 0.05), despite their initial differences (i.e., day 4) (Tukey’s HSD: *p* < 0.001; except between OC_1:1_ and OC_2:1_ communities) (Appendix S2: Figure S3). Here, we also found that algal biomass decreased over time in microcosms with greater sea microbiome dominance (i.e., mixing ratio of 1:2), in contrast to river inoculum-dominated microcosms wherein the biomass increased.

### Community diversity

Out of the identified 461 microeukaryotic OTUs, 84 were shared among all microcosms (Appendix S2: Figure S4). Communities with only sea inoculum (e.g., S and S_x_) had the least numbers of unique OTUs (n_S_ = 5, n_Sx_ = 2). Interestingly, communities exposed to community coalescence harboured the most unique taxa (OC: 15, RC: 23), comprising mostly of fungal taxa. Taxa richness and Shannon’s diversity suggested elevated alpha diversity of microcosms exposed to repeated coalescence (Duncan’s multiple range test for ANOVA: *p* < 0.05), and no significant differences between inoculum sources (except the inverse Simpson’s index) (Appendix S2: Figure S5). Treatments with repeated coalescence and river dominance (e.g., RC_2:1_) had the highest alpha diversity estimates.

### Compositional dynamics imposed by community coalescence

From a compositional point of view, sea inoculum represented a diatom and dinoflagellate-dominated community (Appendix S2: Figure S6), while river inoculum was dominated by golden algae (e.g., Crysophyceae) and ciliates (Appendix S2: Figure S7). Differential abundance analysis revealed ten OTUs with >10 % prevalence (i.e., OTUs present in more than 10% of the samples) in the sea inoculum (mainly Ochrophyta and Dinoflagellata) and forty in the river inoculum (mainly Ochrophyta), that had been negatively affected (*p*_FDR.adj_ < 0.05) by abiotic coalescence (in S_x_ and R_x_) (Appendix S2: Figure S8, S9).

After removing taxa that had been negatively impacted by the environmental coalescence, in order to assess the effect of only biotic community coalescence rather than pure effect of environmental filtering, our differential abundance analysis revealed numerous OTUs (prevalence > 10 %) that decreased or increased in abundance. The relative abundances of these differentially abundant taxonomic groups are presented in Figure 4. We found that mainly diatoms (e.g., *Chaetoceros*, *Thalassiosira* and *Skeletonema*) within the Ochrophyta phylum decreased in abundance after community coalescence (Figure 2). In contrast, community mixing resulted in increased abundances of numerous microeukaryotes including fungi (e.g., Ascomycota, Basidiomycota and early-diverging zoosporic fungi), ciliates and other microeukaryotes (i.e., Cercozoa and Katablepharidophyta) (Figure 2). There were general trends showing that the relative abundances of Cercozoa and chytrids were higher in the final (i.e., day 16) coalesced communities with more sea inoculum (i.e., mixing ratio of 1:2), while Ascomycota, Basidiomycota, Rozellomycota and Chlorophyta OTUs had greater relative abundances in river dominated coalesced communities (i.e., mixing ratio of 2:1).

**Figure 2.**
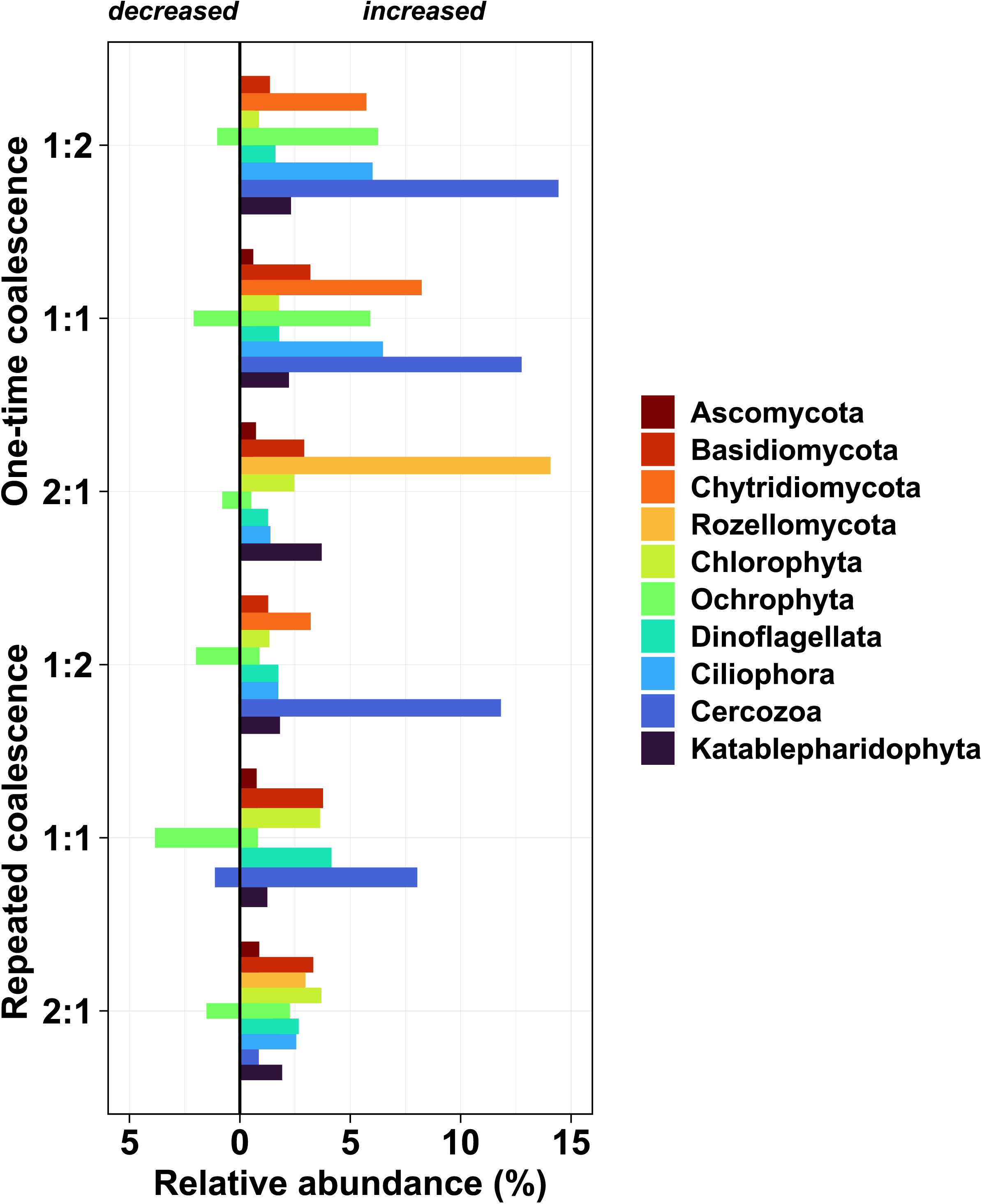
Relative abundances (>0.5 %) of microeukaryotes in the coalesced communities across different community coalescence treatments with three mixing ratios (river:sea). OTUs of coalesced communities showing significant (*p*_FDR.adj_ < 0.05) increase or decrease in taxa abundance, compared to their parent communities, were identified by differential abundance analysis, and grouped by higher taxonomic levels for clarity.

The total relative abundances of the significantly decreased OTUs (selected based on the differential analysis) were significantly higher in repeated versus one-time coalescence treatments (χ^2^ = 23.77, *p* < 0.001). In contrast, coalescence frequency (one-time vs. repeated) had no effect on the relative abundance of OTUs (χ^2^ = 2.19, *p* = 0.139) which maintained or significantly increased (*p*_FDR.adj_ < 0.05), following mixing.

### Community structure and stability

The NMDS of the microeukaryotic communities (Figure 3a) together with the pairwise PERMANOVA results showed that the parent communities (S and R) and those exposed to environmental coalescence (S_x_ and R_x_), by mixing media with 1:1 ratio, were compositionally different (pairwise PERMANOVA, *p* < 0.05), indicating the impact of environmental filtering. Communities exposed to different community coalescence treatments were also significantly different from each other (pairwise PERMANOVA, *p* < 0.05), except for OC_1:1_ and RC_1:1_, as well as OC_1:1_ and RC_1:2_. Although complete convergence to each of the parent community did not occur in any of these communities, the compositions of all community coalesced treatments shifted towards sea parent community (S).

**Figure 3.**
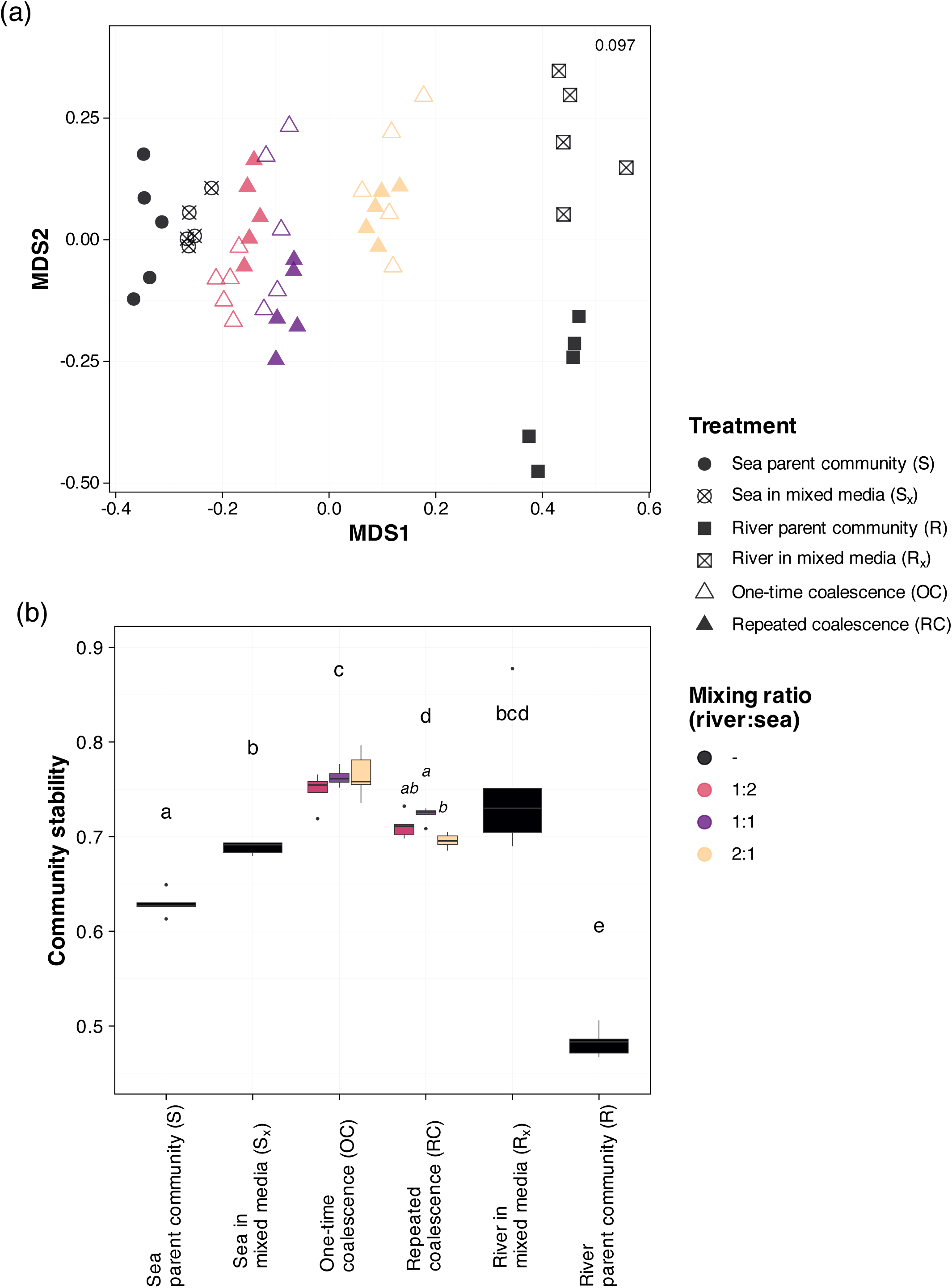
(a) Microeukaryotic community compositions across parent communities and coalescence treatments. Stress value is shown on the upper right corner. (b) Community stabilities based on the ratio of negative:positive cohesion (Herren and McMahon 2017) across parent communities and treatments. Significant (*p* < 0.05) differences in community stability across samples and mixing ratios are represented by lowercase and italicized letters, respectively. N = 5 for each type of sample and treatment. Error bars indicate standard deviations.

When we inferred community stability from the ratio of negative versus positive community cohesion, we found that communities had high proportions of positive cohesion (attributed to facilitative interactions) in all cases (ratio of negative:positive cohesion < |1|), indicating numerous positive-feedback loops that lead to low stability (Herren and McMahon 2017, Hernandez et al. 2021). Nevertheless, sea parent communities (S) were significantly more stable (BH-corrected Wilcoxon test: *p* < 0.05) than river parent communities (R) (Figure 3b), and S community stability increased when they were grown in mixed media (S_x_ and R_x_). Similarly, coalesced communities, OC treatments in particular, had greater community stability (i.e., increased proportion of negative cohesion) than their respective parent communities. Differences in mixing ratios had only an effect in the case of repeated coalescence treatments (Figure 3b), wherein community stability presented significant decreasing difference (BH-corrected Wilcoxon test: *p* < 0.05) between RC_1:1_ and RC_2:1_.

The observed coalesced communities differed in all cases from their corresponding expected community compositions (t-test: *p* < 0.001) and their predictability differed across coalescence treatments (ANOVA: F = 32.23, *p* < 0.0001) (Figure 4a). A clear pattern suggesting decreasing predictability of the coalesced communities with increasing ratio of river inoculum (i.e., predictability of river:sea mixing: 2:1 < 1:1 < 1:2) was found (Figure 4a). In addition, predictability was typically lower in communities exposed to one-time coalescence than in those subjected to repeated coalescence (*p* < 0.05).

**Figure 4.**
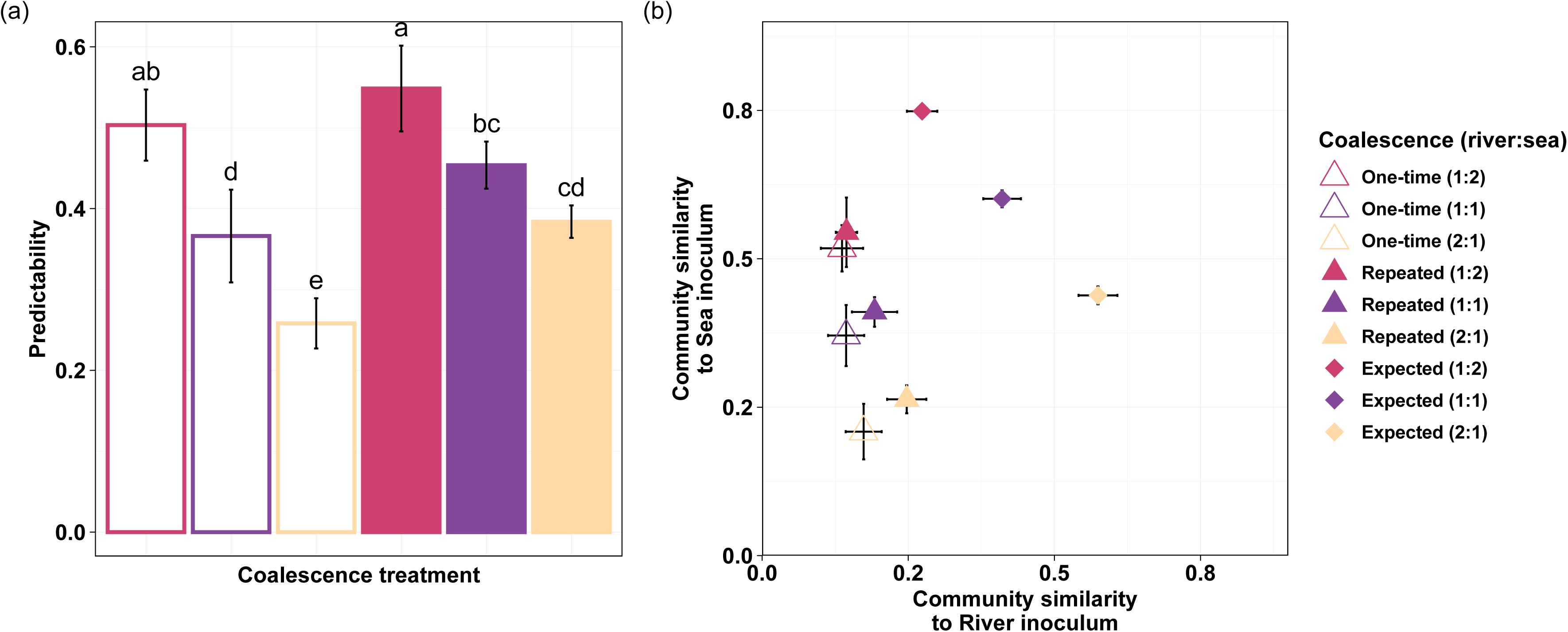
(a) Predictability of community coalescence outcomes increases with the ratio of sea community in the final coalesced communities based on the similarity of the observed and expected communities. All observed communities significantly differ from the expected communities (*p* < 0.001). Significant (*p* < 0.05) differences among treatments are represented by lowercase letters. (b) Similarity of the observed and expected coalesced communities in relation to their parent communities. Expected community similarities were determined by conservative mixing model based on the applied mixing ratio or river and sea parent community (see Methods for details). N = 5 for each type of treatment. Error bars indicate standard deviations.

The observed Bray-Curtis similarity between each coalesced community and its parent community was calculated and compared to the expected similarity (Figure 4b). This showed that all communities had lower similarity to their parent communities than expected (ANOVA: F = 69.12, p < 0.001) (Figure 4b). Furthermore, communities that were either exposed to one-time coalescence treatment (i.e., empty triangles) or received higher proportion of river inoculum (i.e., yellow triangles) tended to diverge even more from the expected similarities than those exposed to repeated coalescence treatments and having lower ratio of river inoculum, respectively. In general, coalesced communities showed greater similarity to sea than to river parent communities (Figure 4b).

## Discussion

In this study we mimicked estuarine conditions by mixing a sub-arctic freshwater river community with brackish water community of the Gulf of Bothnia and assessed the outcome of different mixing scenarios. We observed asymmetrical community coalescence outcome as coalesced communities were generally shifted towards the sea parent community which was also generally less impacted by the effect of environmental coalescence (i.e., environmental filtering). Community coalescence increased community stability and most likely promoted competitive interactions with the introduced species, leading to a stabilizing effect by negative-feedback loops. Overall, predictability of coalescence outcomes was greater when the initially more stable fraction (i.e., sea microbes) dominated the final community and this predictability increased when communities were repeatedly mixed.

### Compositional dynamics imposed by coalescences

The sampled water bodies are characterized by oligotrophic conditions (Andersson et al. 1996, Wasmund et al. 2001, Wikner and Andersson 2012). Specifically, our inocula originated from phosphorus- and nitrogen-limited river and sea habitats, respectively. In such oligotrophic environments we expect species to be under higher stress than in nutrient-rich environments (Ornolfsdottir 2004).

River communities subjected to environmental coalescence suffered from a four times greater taxa loss (8 % of riverine OTUs), compared to sea microbiome (2 % of marine OTUs). This suggests, in line with Cloern et al. (2017) and Rocca et al. (2020), that sea microeukaryotes are better adapted to the new environment imposed by habitat mixing, probably due to their brackish origin, making them more tolerant to saline conditions than freshwater species. However, most diatoms, the group which suffered the most from the biotic effects of community coalescence, originated from sea communities.

In community coalescence treatments, unequal mixing ratios of river and sea communities resulted in contrasting algal biomass, wherein primary producers decreased in sea-dominated coalesced communities, while increased in river-dominated microcosms over time. A possible explanation for this phenomenon is that the more saline mixed medium causes riverine algal biomass to decline (i.e., filtered by the environment), which leads to the opening of niches that the more salt tolerant sea alga can occupy. Nevertheless, it seems that changes in water conditions are less influential for microeukaryotes in general, as evidenced by the low percentages of species loss during environmental coalescences, than that of bacterial communities exposed to various salinity disturbances (Shen et al. 2018, Bier et al. 2022). This might be the consequence of the small changes in salinity within the tolerance range of microeukaryotes, or due to complex ecological interactions where changes in salinity due to habitat mixing influence heterotrophic microeukaryotes. For example, the decline of certain microbes, as discussed above, opens niches and supplies organic matter for the microbial loop, supporting bacterial growth and thereby bactivorous microeukaryotes (Stefanidou et al. 2018). We can speculate that this process can further be promoted by the elevated photosynthesis (observed in river-dominated coalesced communities, see e.g., Appendix S2: Figure S3) that might have increased pH (not measured herein) and in turn released coprecipitated P into the water, as evidenced by the increased TDP levels in coalesced communities. In addition to environmental filtering, enhanced biotic interactions (e.g., diatom – chytrids) may have played a relevant role in the in sea-dominated coalesced communities (Vass et al. 2022), contributing to the species loss and the observed trend of declining biomass of primary producers (e.g., diatoms) in sea-dominated coalesced microcosms (i.e., RC_1:2_/OC_1:2_), while elevating the abundance of chytrids (parasitic zoosporic fungi).

The high number of unique microeukaryotes in coalesced communities suggests and supports earlier finding that rare microbial taxa emerge during mixing events (Rocca et al. 2020). Such phenomena are most likely attributed to the selective advantage of certain phenotypes of these microbes under the new coalesced conditions, as well as the earlier described potential decline of certain microbes that opens niches and supports the establishment of emerging taxa. This explanation is also in line with the increased community stability (i.e., elevated fraction of competitive interactions) observed in the environmental coalescence treatments.

### Coalescence mediates community stability

Community stability can be inferred from numerous community properties (Shade et al. 2012). Here, we approached community stability from the point of community cohesion, a metric that estimates the connectivity of microbial communities that stems from biotic associations (Herren and McMahon 2017). As the authors highlight, taxa associations arise from biotic interactions and environmental drivers. Thanks to the environmental coalescence treatments, we were able to rule out the effects of environmental filtering on coalesced communities, hence, gaining a stronger support for competitive interactions alone when negative cohesion emerged. Positive cohesion can be indicative for both facilitative interactions and environmental synchrony and these two cannot be disentangled in our present study. Nevertheless, the ratio of negative and positive cohesion allowed us to infer community stability of our observed communities, given that the cohesion values are indicative of negative- and positive-feedback loops, promoting or reducing community stability, respectively (Mitri and Richard Foster 2013, Coyte et al. 2015, Herren and McMahon 2017).

Although our findings suggest greater overall dominance of facilitative interactions and/or the influence of environmental synchrony (that is, the dominance of positive cohesion) across treatments, such dominance was limited by coalescence treatments, elevating the importance of competitive interactions that tend to be more evidenced in microbial communities (Foster and Bell 2012). The weakened dominance of facilitative interactions could potentially be attributed to disappearing reciprocal benefits (e.g., metabolic cross-feeding) since spatial structuring, that has similar effect, are unlikely in our microcosms (Harcombe 2010). In such scenario, the importance of ecological co-selection, a phenomenon which aids members of a community to recruit one another, can be diminished and dominant taxa could not invade another community on their own, successfully (Diaz-Colunga et al. 2022). Although this reasoning is experimentally not tested herein, the increased levels of the inverse Simpson’s index in repeatedly coalesced communities (see e.g., Appendix S2: Figure S5) may suggest such phenomenon as it indicates mechanisms that counteract dominance. This might also explain why numerous microeukaryotes such as ciliates and parasitic fungi could elevate their abundances, following coalescence. This and other processes generated by environmental coalescence could have provided avenues for the introduction of additional species and/or the emergence of rare taxa that triggered competition to a greater extent and by doing so, leading communities towards greater stability. The driving mechanism behind this, as introduced earlier, originates from the increased number of negative-feedback loops which dampen the destabilizing effect of facilitative interactions that would otherwise lead to species loss. Our diversity estimates can support this reasoning as taxa richness increased in the coalesced communities, particularly in those that experienced repeated coalescence events, adding further evidence for a species maximum of microeukaryotes in brackish conditions (Filker et al. 2019, Tee et al. 2021).

Overall, our findings that community coalescence in estuaries enhances microeukaryotic community stability by increasing the number of competitions supports the notion that dampening the proportion of positive-feedback loops leads to even greater stability in microbial communities (Coyte et al. 2015), but questions May’s (1972) and Coyte et al.’s (2015) work on the destabilizing effect of increasing species diversity.

The composition of the coalesced communities with greater mixing ratio of the more stable sea parent community generated greater predictabilities, suggesting that greater community stability can be associated with better predictability of community coalescence outcomes. This likelihood can be further enhanced as the frequency of mixing increases (i.e., repeatedly colliding communities). Lechón-Alonso et al.’s (2021) simulation study suggested that communities experiencing repeated mixing events should gradually shift from competitive towards more facilitative communities, which we did not find support for in this 16-day long study. Instead, regardless of the frequency of coalescence events (i.e., one-time vs. repeated), our coalesced communities became more competitive than their parent communities.

## Conclusion

Our findings suggest that the parent community with greater stability determines coalescence outcome, where the fate of its members is influenced by the mixing proportion of parent communities and to a lesser extent the temporal dynamics of community coalescence (one-time versus regular exchange). Nevertheless, our finding that community coalescence increases microeukaryotic diversity and promotes stability should be tested on microbial communities originating from other climatic regions and estuary systems with greater differences in salinity between endmembers to see how these results can be generalized across estuaries.

Understanding the outcomes of community coalescence and the fate of microbes in their mixed environment are essential to understand and model biodiversity and associated functionality. A changed climate could modify coalescence outcomes (Vass et al. 2021), and even variation in weather conditions trigger more frequent and more intense mixing scenarios (e.g., flooding and soil runoff into streams/rivers in response to heavy rainfalls) (Mansour et al. 2018). These processes will inevitably impact all features of estuarine ecosystems, including diversity, composition, function, and its capability to respond to various disturbances (Rocca et al. 2021).

## Acknowledgements

We are grateful for the support of Umeå Marine Science Centre for collecting offshore sample and performing chemical measurements. We are especially thankful to Fernanda Helena Bosco De Miranda Vasconcelos for providing the AquaPen device and assistance. Funding for this project was provided by Umeå University and the Swedish strategic marine research programme EcoChange (Swedish Research Council Formas). This research was conducted using the resources of High-Performance Computing Center North (HPC2N) and Uppsala Multidisciplinary Center for Advanced Computational Science (UPPMAX). The computations and data handling were enabled by resources provided by the Swedish National Infrastructure for Computing (SNIC), partially funded by the Swedish Research Council through grant agreement no. 2018-05973.

## Author contributions

AA, UCG, JW conceived the study and obtained funding for this research. AA and MV conducted sample collection. MV developed the research questions, performed sample processing and the experiment, processed samples, analysed the data. MV drafted the manuscript with substantial help from ASz. All authors contributed substantially to interpretation of results and writing.

