## Appendix S2 for "Community stability increases the predictability of microeukaryote community coalescence outcomes"

### Community stability increases the predictability of microeukaryote community coalescence outcomes

*Journal: Ecology*

**Máté Vass[1,2*], Anna Székely[3], Ulla Carlsson-Graner[1], Johan Wikner[1,4], Agneta Andersson[1,4]**

[1]Department of Ecology and Environmental Science, Umeå University, SE-901 87, Umeå, Sweden
[2]Division of Systems and Synthetic Biology, Department of Life Sciences, Science for Life Laboratory, Chalmers University of Technology, SE-412 96, Gothenburg, Sweden
[3]Department of Aquatic Sciences and Assessment; Division of Microbial Ecology, Swedish University of Agricultural Sciences, SE-750 07, Uppsala, Sweden
[4]Umeå Marine Sciences Centre, Umeå University, SE-905 71, Hörnefors, Sweden

*corresponding author's current address:, Division of Systems and Synthetic Biology, Department of Life Sciences, Science for Life Laboratory, Chalmers University of Technology, SE-412 96, Gothenburg, Sweden

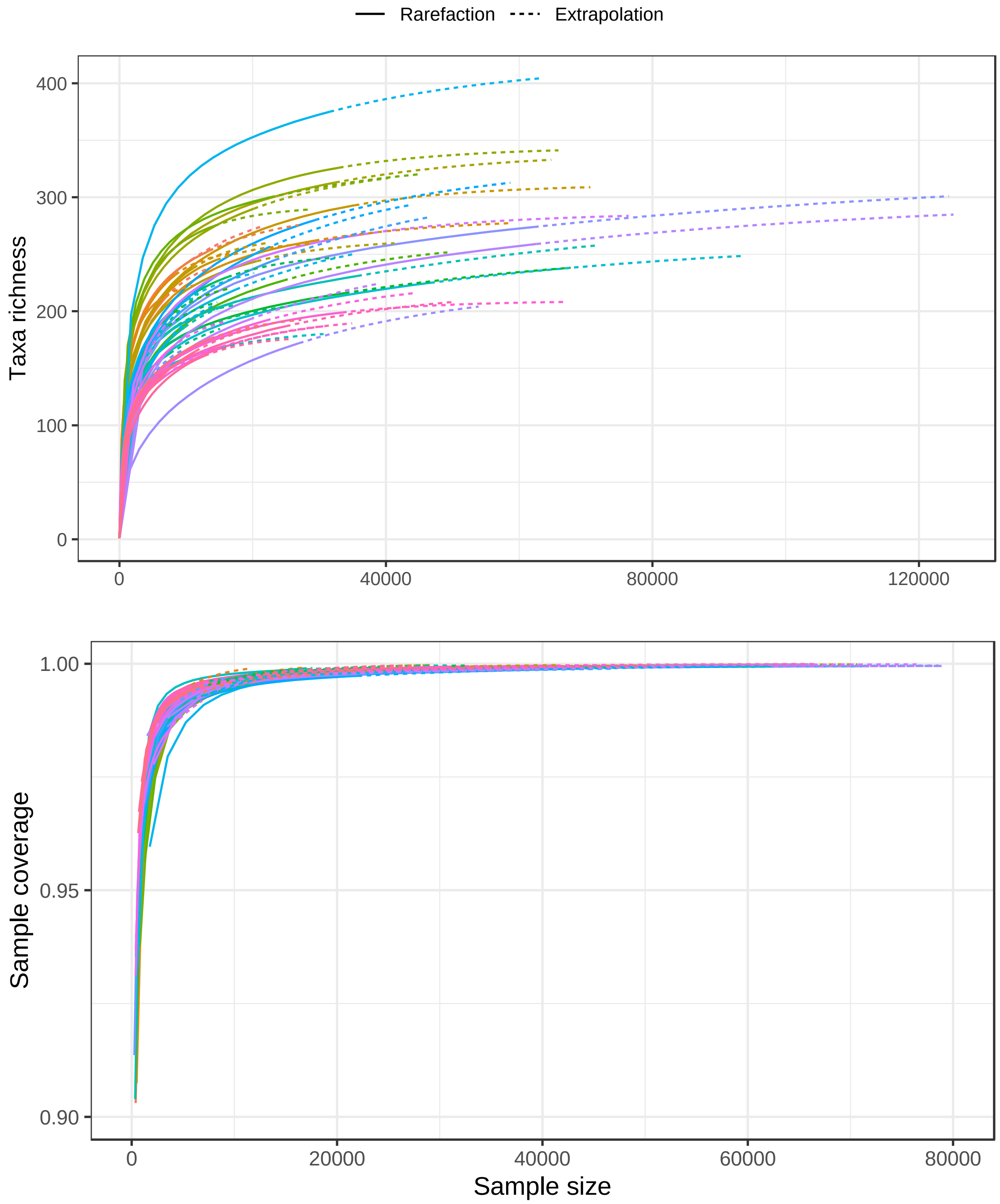


**Figure S1.** Diversity estimates and sample completeness (coverage) curves. Colours indicate different samples.

**Equation S1: Conservative mixing model**

We calculated the proportion of OTUs in the expected coalesced communities (ECC) based on the OTU proportions in the two parent communities (R and S) and the applied mixing ratios (i.e., 1:1, 1:2 or 2:1):

$$ECC = \sum_{i=1}^{n}(RA_{R,i} * MR_R + RA_{S,i} * MR_S)$$

Where $i$ is an OTU, $n$ is the total number of OTUs in the communities, $RA_{R,i}$ and $RA_{S,i}$ are the relative abundance of $i$ OTU in parent community R and S, respectively, and $MR_R$ and $MR_S$ are the mixing ratio of R and S, respectively.

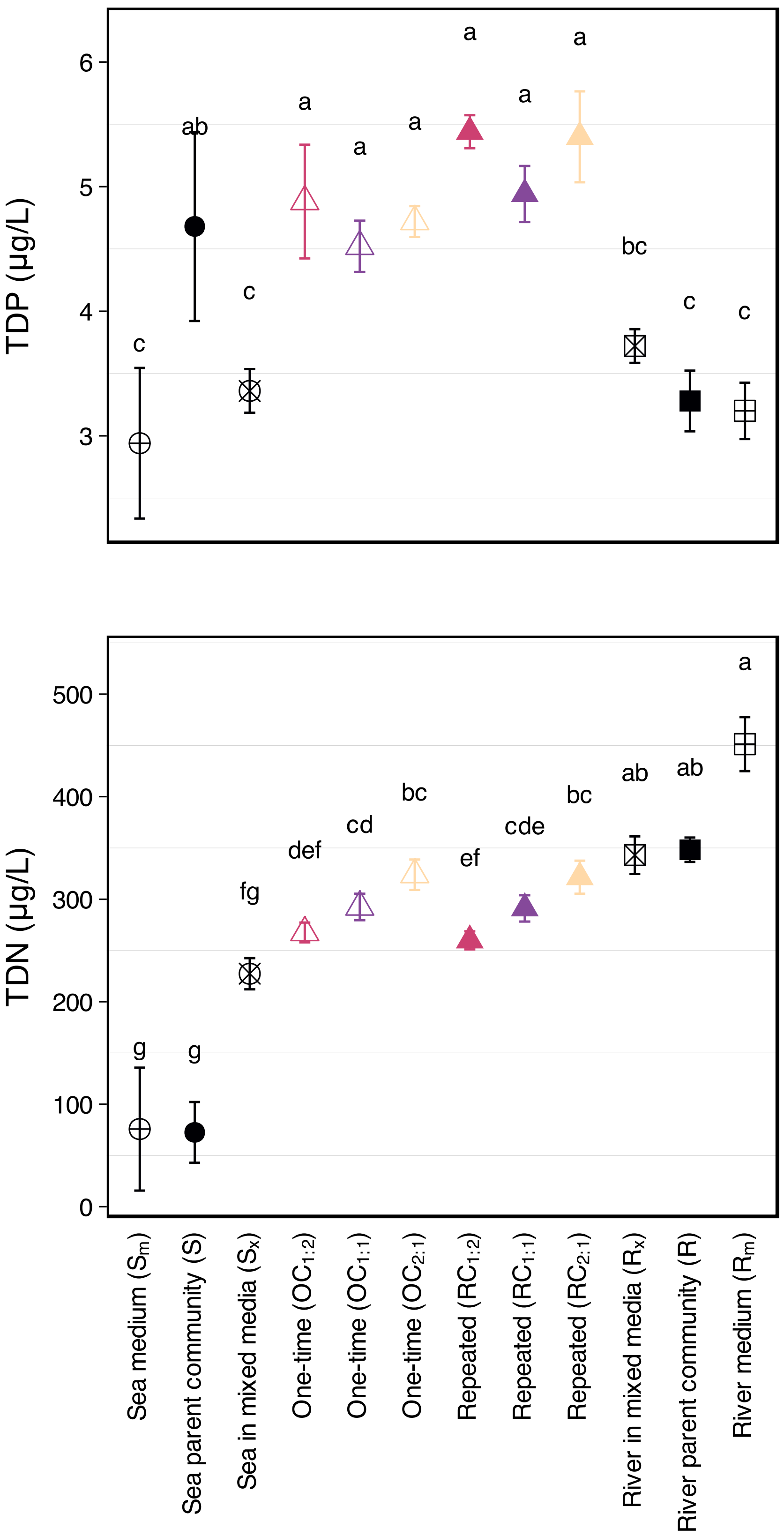


**Figure S2**. Differences in total dissolved phosphorus (TDP) and total dissolved nitrogen (TDN) across inoculum sources and treatments. N = 5 for each type of treatment. Significant (Kruskal-Wallis; $p_{adj} < 0.05$) differences are represented by lowercase letters. Error bars indicate standard deviations, and parentheses represent the mixing ratio of river:sea of coalesced communities.

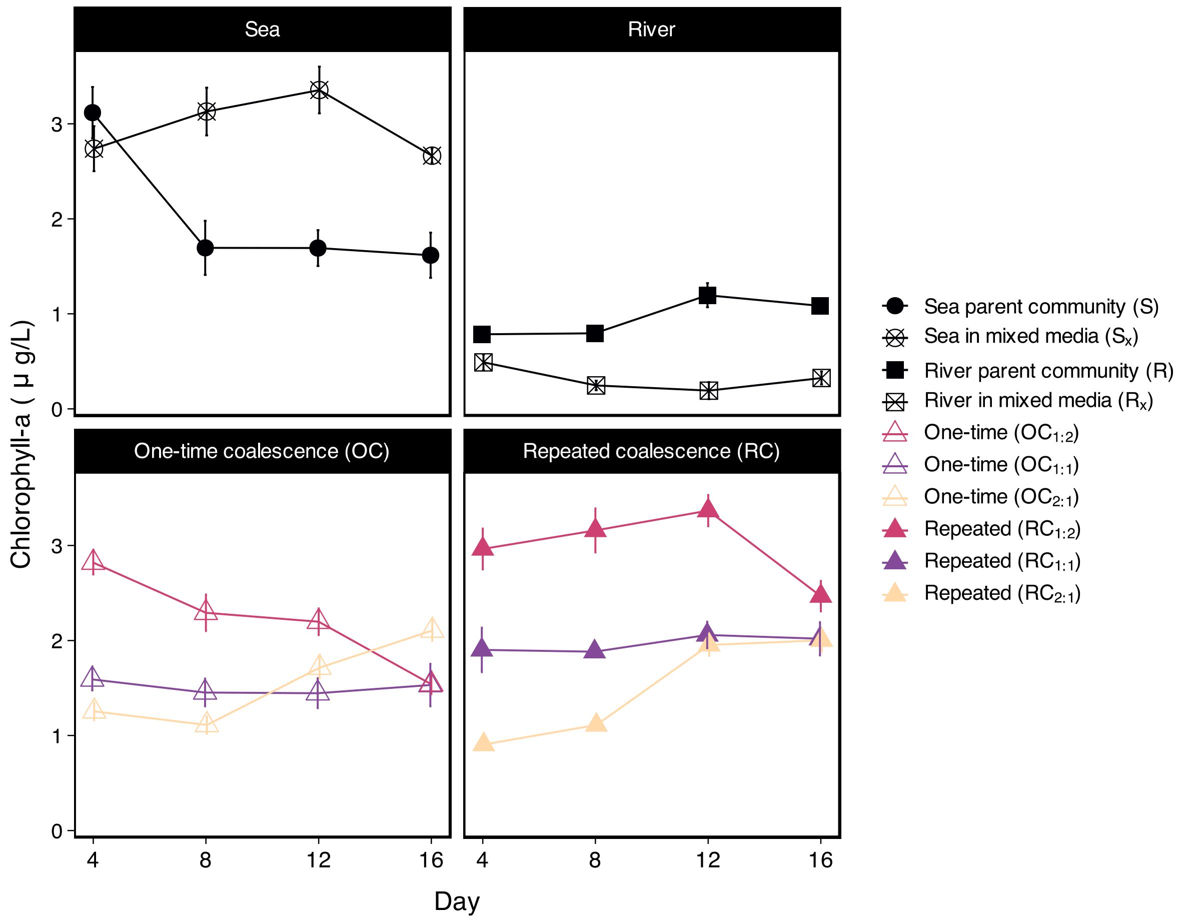


**Figure S3**. Changes in chlorophyll-a (as an estimator of the biomass of primary producers) as estimated by measuring the chlorophyll fluorescence-induced dynamic curve using AquaPen-C device (Photon Systems Instruments, Brno, Czechia). N = 5 for each type of culture and treatment at each date. Error bars indicate standard deviations, and parentheses represent the mixing ratio of river:sea of coalesced communities.

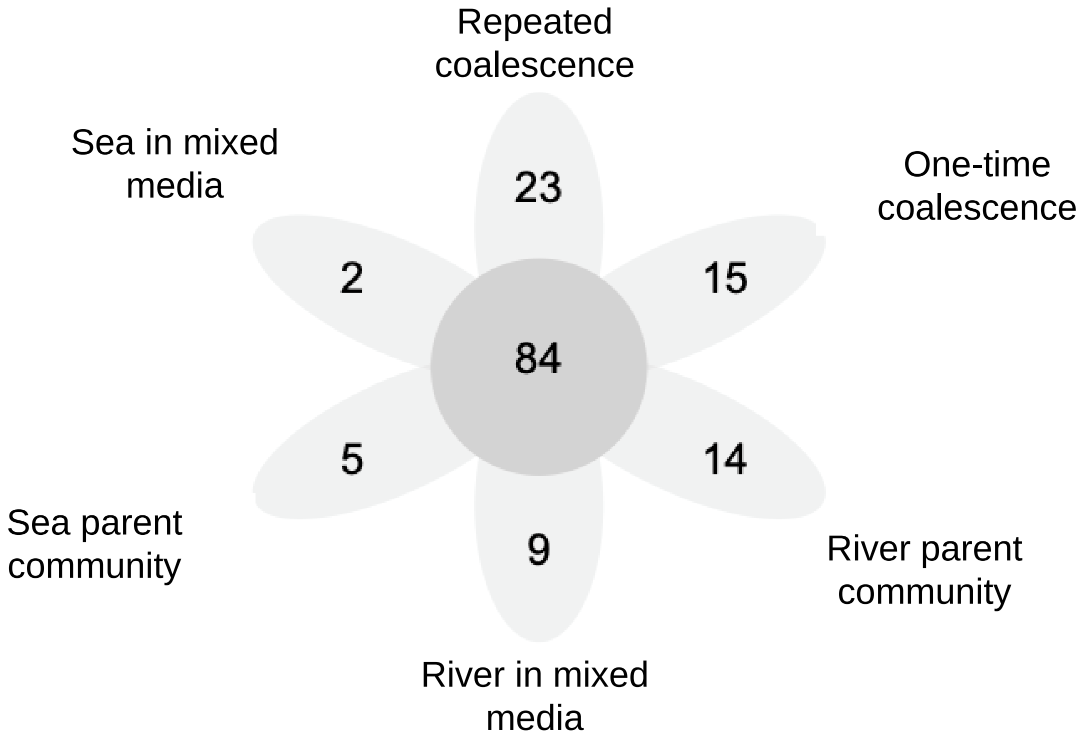


**Figure S4**. Number of unique and shared OTUs in the different inoculum sources and treatments. N = 5 for each type of sample and treatment.

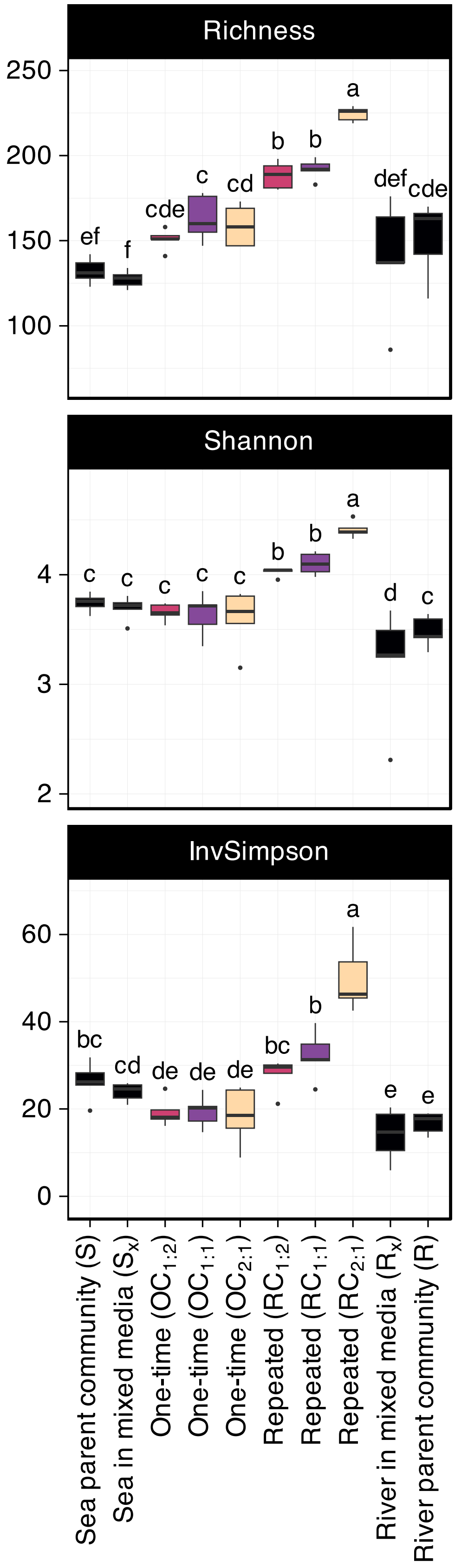


**Figure S5.** Alpha diversity estimates across samples and treatments. Significant post-hoc groups (Duncan's multiple range test for ANOVA; $p < 0.05$) are represented by lowercase letters. N = 5 for each type of each treatment. Error bars indicate standard deviations, and the mixing ratios of one-time and repeated treatments are indicated by the ratio of river:sea.

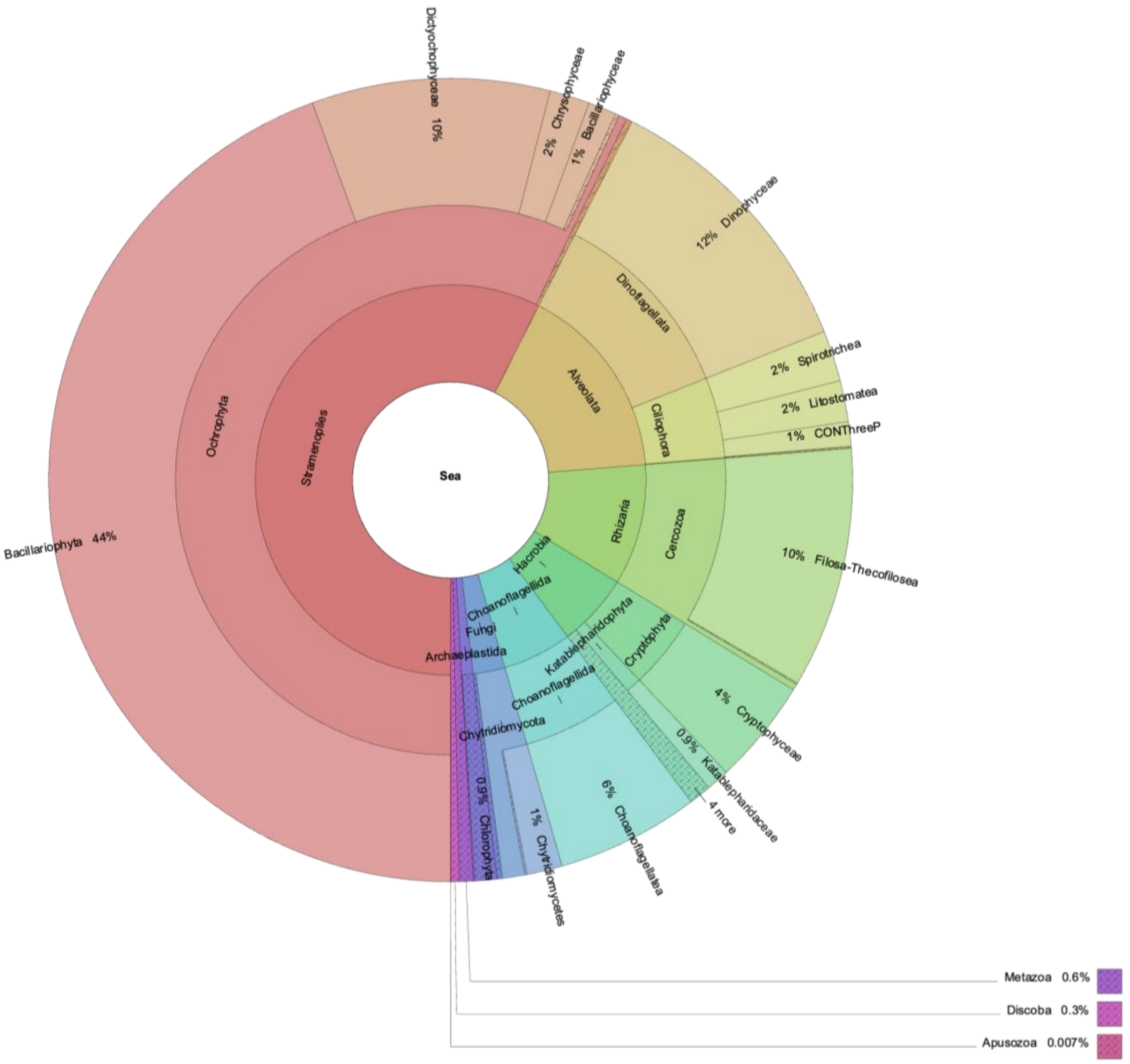


**Figure S6**. Taxonomic distributions of microeukaryotes, indicated as percentages of the total number of reads, found (day 16) in the sea inoculum microcosms (N = 5).

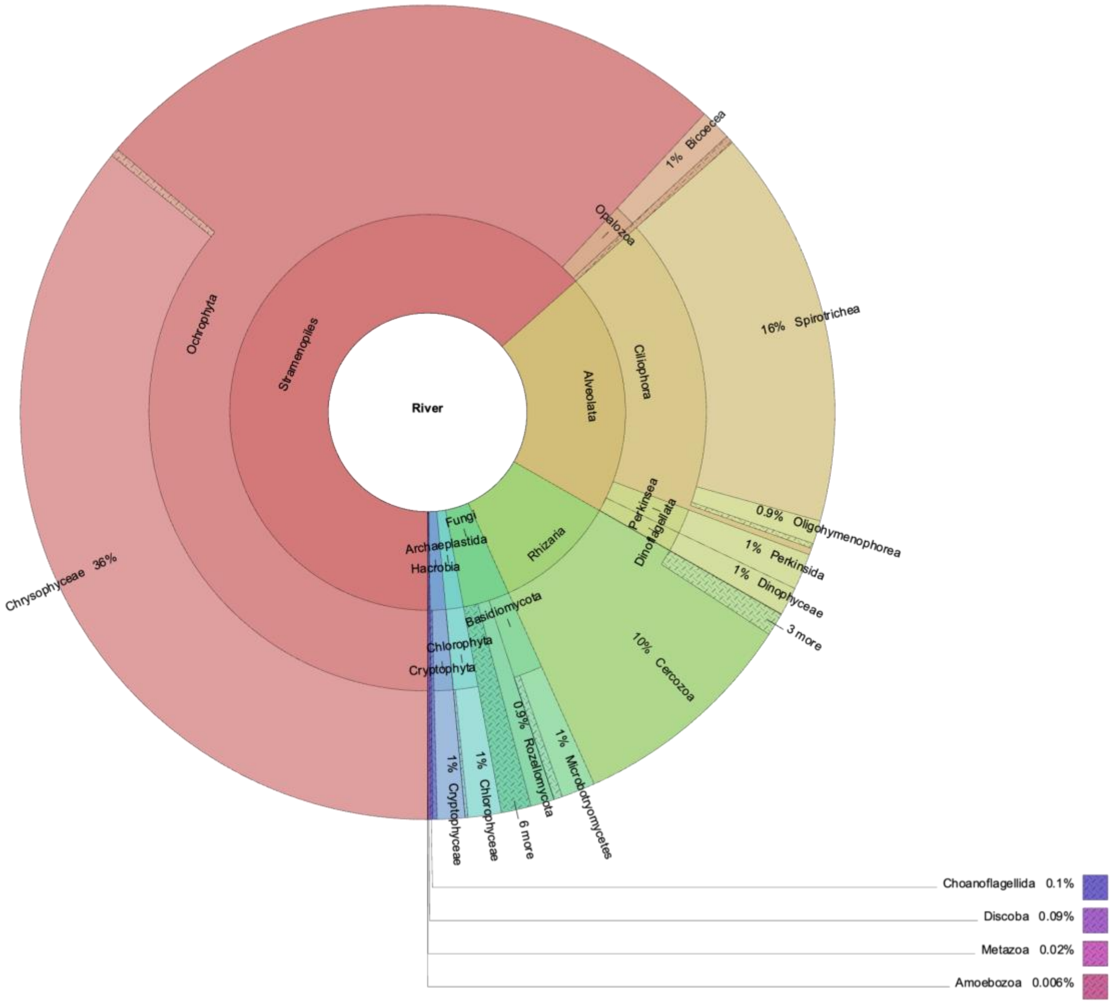


**Figure S7**. Taxonomic distributions of microeukaryotes, indicated as percentages of the total number of reads, found (day 16) in the river inoculum microcosms (N = 5).

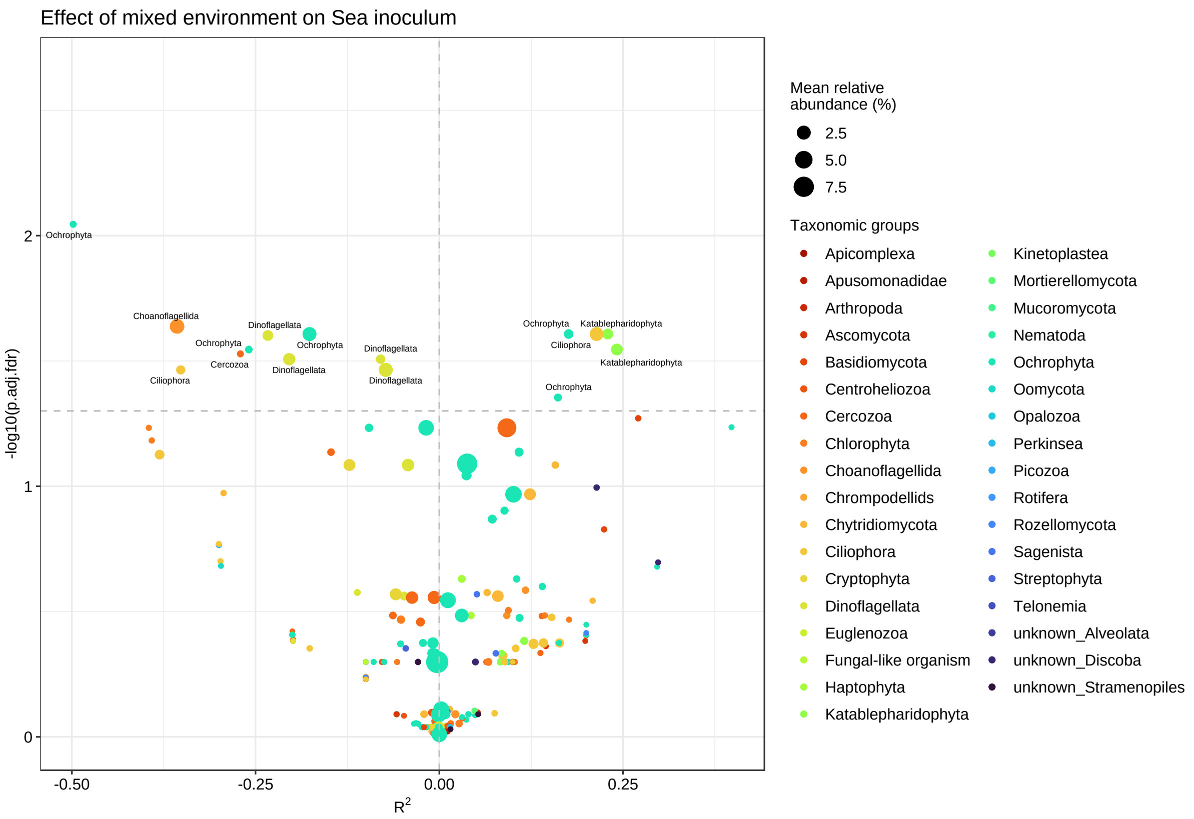


**Figure S8.** Differential abundance analyses of microeukaryotes of sea parent community after being incubated in the mixed environment (1:1 mixture of sea and river medium). OTUs above the horizontal dashed line indicate significant (*p* < 0.05; permutation-based FDR-adjusted p-values) increase ($R^2 > 0$) or decrease ($R^2 < 0$) in taxa abundance as assessed by ZicoSeq (Yang & Chen 2022).

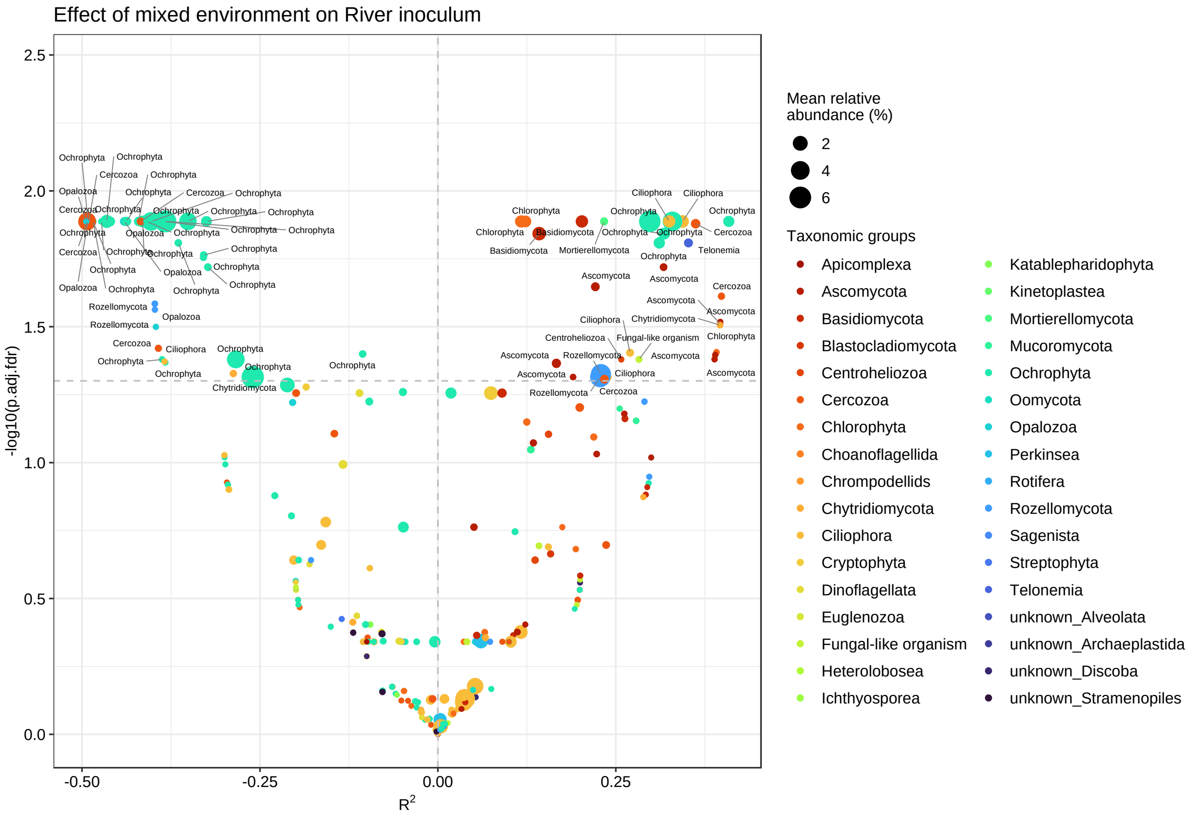


**Figure S9.** Differential abundance analyses of microeukaryotes of river parent community after being incubated in the mixed environment (1:1 mixture of sea and river medium). OTUs above the horizontal dashed line indicate significant (*p* < 0.05; permutation-based FDR-adjusted p-values) increase ($R^2 > 0$) or decrease ($R^2 < 0$) in taxa abundance as assessed by ZicoSeq (Yang & Chen 2022).
