## Appendix S1 for "Community stability increases the predictability of microeukaryote community coalescence outcomes"

**Community stability increases the predictability of microeukaryote community coalescence outcomes**

*Journal: Ecology*

**Máté Vass[1,2*], Anna Székely[3], Ulla Carlsson-Graner[1], Johan Wikner[1,4], Agneta Andersson[1,4]**

[1]Department of Ecology and Environmental Science, Umeå University, SE-901 87, Umeå, Sweden
[2]Division of Systems and Synthetic Biology, Department of Life Sciences, Science for Life Laboratory, Chalmers University of Technology, SE-412 96, Gothenburg, Sweden
[3]Department of Aquatic Sciences and Assessment; Division of Microbial Ecology, Swedish University of Agricultural Sciences, SE-750 07, Uppsala, Sweden
[4]Umeå Marine Sciences Centre, Umeå University, SE-905 71, Hörnefors, Sweden

*corresponding author's current address:, Division of Systems and Synthetic Biology, Department of Life Sciences, Science for Life Laboratory, Chalmers University of Technology, SE-412 96, Gothenburg, Sweden

**Table S1. Barcoded primers used in the study**

| name | index | index | primer | name | fused_sequence | Tm | checked: http://biotools.nubic.northwestern.edu/OligoCalc.html |
|---|---|---|---|---|---|---|---|
| Balzano_F | 1 | ACAGACGACTACAAACGGAATCGA | CCAGCASCYGCGGTAATTCC | Balzano_F_1 | ACAGACGACTACAAACGGAATCGACCAGCASCYGCGGTAATTCC | 71 | good |
| Balzano_F | 2 | TCGATTCCGTTTGTAGTCGTCTGT | CCAGCASCYGCGGTAATTCC | Balzano_F_2 | TCGATTCCGTTTGTAGTCGTCTGTCCAGCASCYGCGGTAATTCC | 71 | good |
| Balzano_F | 3 | AACTAGGCACAGCGAGTCTTGGTT | CCAGCASCYGCGGTAATTCC | Balzano_F_3 | AACTAGGCACAGCGAGTCTTGGTTCCAGCASCYGCGGTAATTCC | 72 | good |
| Balzano_F | 4 | TTCGGATTCTATCGTGTTTCCCTA | CCAGCASCYGCGGTAATTCC | Balzano_F_4 | TTCGGATTCTATCGTGTTTCCCTACCAGCASCYGCGGTAATTCC | 70 | good |
| Balzano_F | 5 | GAGAGGACAAAGGTTTCAACGCTT | CCAGCASCYGCGGTAATTCC | Balzano_F_5 | GAGAGGACAAAGGTTTCAACGCTTCCAGCASCYGCGGTAATTCC | 71 | good |
| Balzano_F | 6 | ATCTCTTGACACTGCACGAGGAAC | CCAGCASCYGCGGTAATTCC | Balzano_F_6 | ATCTCTTGACACTGCACGAGGAACCCAGCASCYGCGGTAATTCC | 72 | good |
| Balzano_F | 7 | ATGAGTTCTCGTAACAGGACGCAA | CCAGCASCYGCGGTAATTCC | Balzano_F_7 | ATGAGTTCTCGTAACAGGACGCAACCAGCASCYGCGGTAATTCC | 71 | good |
| 3143R | 1 | ACCGAGATCCTACGAATGGAGTGT | RCCACAAGCYARTTATCC | 3143R_1 | ACCGAGATCCTACGAATGGAGTGTRCCACAAGCYARTTATCC | 67 | good |
| 3143R | 2 | ACGGTATGTCGAGTTCCAGGACTA | RCCACAAGCYARTTATCC | 3143R_2 | ACGGTATGTCGAGTTCCAGGACTARCCACAAGCYARTTATCC | 67 | good |
| 3143R | 3 | AGGTGATCCCAACAAGCGTAAGTA | RCCACAAGCYARTTATCC | 3143R_3 | AGGTGATCCCAACAAGCGTAAGTARCCACAAGCYARTTATCC | 66 | good |
| 3143R | 4 | TTCTGAAGTTCCTGGGTCTTGAAC | RCCACAAGCYARTTATCC | 3143R_4 | TTCTGAAGTTCCTGGGTCTTGAACRCCACAAGCYARTTATCC | 66 | ok |
| 3143R | 5 | GACAGACACCGTTCATCGACTTTC | RCCACAAGCYARTTATCC | 3143R_5 | GACAGACACCGTTCATCGACTTTCRCCACAAGCYARTTATCC | 67 | good |
| 3143R | 6 | TTCTCAGTCTTCCTCCAGACAAGG | RCCACAAGCYARTTATCC | 3143R_6 | TTCTCAGTCTTCCTCCAGACAAGGRCCACAAGCYARTTATCC | 67 | ok |
| 3143R | 7 | TAGCTGACTGTCTTCCATACCGAC | RCCACAAGCYARTTATCC | 3143R_7 | TAGCTGACTGTCTTCCATACCGACRCCACAAGCYARTTATCC | 67 | good |
| 3143R | 8 | TACAAGCATCCCAACACTTCCACT | RCCACAAGCYARTTATCC | 3143R_8 | TACAAGCATCCCAACACTTCCACTRCCACAAGCYARTTATCC | 66 | good |

**Table S2. Summary of sequence processing**

| Raw good quality reads (Q>9) before demultiplexing | Fasta_file_name | Sample_ID | Raw good quality reads (Q>9) | Filtered reads (2-6 kbp) | Final reads used for consensus creation | Coverage (%) |
|---|---|---|---|---|---|---|
| **2 436 290** | Euk_F11A.fastq | F11A | 19693 | 18932 | 18449 | **93.68** |
| | Euk_F11B.fastq | F11B | 10680 | 10161 | 9873 | **92.44** |
| | Euk_F11C.fastq | F11C | 21735 | 20764 | 20222 | **93.04** |
| | Euk_F11D.fastq | F11D | 8084 | 7678 | 7446 | **92.11** |
| | Euk_F11E.fastq | F11E | 12794 | 12278 | 11982 | **93.65** |
| | Euk_F12A.fastq | F12A | 44856 | 42583 | 41029 | **91.47** |
| | Euk_F12B.fastq | F12B | 24285 | 22920 | 22037 | **90.74** |
| | Euk_F12C.fastq | F12C | 44526 | 42282 | 40808 | **91.65** |
| | Euk_F12D.fastq | F12D | 16516 | 15538 | 15045 | **91.09** |
| | Euk_F12E.fastq | F12E | 35955 | 34316 | 33276 | **92.55** |
| | Euk_F21A.fastq | F21A | 57259 | 55312 | 54401 | **95.01** |
| | Euk_F21B.fastq | F21B | 29339 | 28357 | 27817 | **94.81** |
| | Euk_F21C.fastq | F21C | 53050 | 51156 | 50412 | **95.03** |
| | Euk_F21D.fastq | F21D | 22595 | 21557 | 21223 | **93.93** |
| | Euk_F21E.fastq | F21E | 49501 | 48140 | 47630 | **96.22** |
| | Euk_O11A.fastq | O11A | 51396 | 49168 | 48297 | **93.97** |
| | Euk_O11B.fastq | O11B | 24113 | 23002 | 22568 | **93.59** |
| | Euk_O11C.fastq | O11C | 54810 | 52091 | 51075 | **93.19** |
| | Euk_O11D.fastq | O11D | 19398 | 18435 | 18123 | **93.43** |
| | Euk_O11E.fastq | O11E | 45960 | 44215 | 43545 | **94.75** |
| | Euk_O12A.fastq | O12A | 29592 | 28271 | 27672 | **93.51** |
| | Euk_O12B.fastq | O12B | 18231 | 17174 | 16706 | **91.64** |
| | Euk_O12C.fastq | O12C | 22189 | 20823 | 20258 | **91.30** |
| | Euk_O12D.fastq | O12D | 11917 | 11211 | 10901 | **91.47** |
| | Euk_O12E.fastq | O12E | 16359 | 15176 | 14838 | **90.70** |
| | Euk_O21A.fastq | O21A | 58141 | 55141 | 54218 | **93.25** |
| | Euk_O21B.fastq | O21B | 28811 | 27504 | 27055 | **93.91** |
| | Euk_O21C.fastq | O21C | 72461 | 70022 | 69209 | **95.51** |
| | Euk_O21D.fastq | O21D | 21035 | 19835 | 19597 | **93.16** |
| | Euk_O21E.fastq | O21E | 38110 | 36355 | 36090 | **94.70** |
| | Euk_R0.fastq | R0 | 41171 | 37798 | 36461 | **88.56** |
| | Euk_RA.fastq | RA | 11067 | 9109 | 8936 | **80.74** |
| | Euk_RB.fastq | RB | 27308 | 23737 | 22959 | **84.07** |
| | Euk_RC.fastq | RC | 36899 | 32108 | 31223 | **84.62** |
| | Euk_RD.fastq | RD | 28600 | 25808 | 24905 | **87.08** |
| | Euk_RE.fastq | RE | 7728 | 5777 | 5685 | **73.56** |
| | Euk_RxA.fastq | RxA | 80653 | 66804 | 66437 | **82.37** |
| | Euk_RxB.fastq | RxB | 29520 | 28369 | 28263 | **95.74** |
| | Euk_RxC.fastq | RxC | 73147 | 68181 | 67495 | **92.27** |
| | Euk_RxD.fastq | RxD | 23101 | 21150 | 20979 | **90.81** |
| | Euk_RxE.fastq | RxE | 51830 | 47629 | 47000 | **90.68** |
| | Euk_S0.fastq | S0 | 5063 | 4144 | 3984 | **78.69** |
| | Euk_SA.fastq | SA | 10679 | 9549 | 9084 | **85.06** |
| | Euk_SB.fastq | SB | 10572 | 10014 | 9511 | **89.96** |
| | Euk_SC.fastq | SC | 27381 | 25713 | 24138 | **88.16** |
| | Euk_SD.fastq | SD | 45436 | 42825 | 40486 | **89.11** |
| | Euk_SE.fastq | SE | 25532 | 24280 | 23273 | **91.15** |
| | Euk_SxA.fastq | SxA | 32968 | 31563 | 30402 | **92.22** |
| | Euk_SxB.fastq | SxB | 47036 | 44965 | 43099 | **91.63** |
| | Euk_SxC.fastq | SxC | 26860 | 25882 | 24881 | **92.63** |
| | Euk_SxD.fastq | SxD | 18849 | 17862 | 17063 | **90.52** |
| | Euk_SxE.fastq | SxE | 26388 | 25035 | 23973 | **90.85** |
| | | **Mean:** | **31 753** | **29 783** | **29 078** | **90.8** |
| | | **Min:** | **5 063** | **4 144** | **3 984** | **73.6** |
| | | **Max:** | **80 653** | **70 022** | **69 209** | **96.2** |
| | | **Total:** | **1 651 179** | **1 548 699** | **1 512 039** | |

Retained seqs: 62.1%
